# Growth instabilities shape morphology and genetic diversity of cellular aggregates

**DOI:** 10.1101/2022.03.28.486080

**Authors:** Alexander Golden, Ilija Dukovski, Daniel Segrè, Kirill S. Korolev

## Abstract

Cellular aggregates assume an incredible variety of shapes ranging from circular molds to irregular tumors. While we understand many of the mechanisms responsible for these spatial patterns, little is known about how the shape of an aggregate influences its ecology and evolution. Here, we investigate this relationship in the context of microbial colonies grown on hard agar plates. This a well-studied system that exhibits a transition from smooth circular disks to more irregular and rugged shapes as either the nutrient concentration or cellular motility is decreased. Starting from a mechanistic model of colony growth, we identify two dimensionless quantities that determine how morphology and genetic diversity of the population depend on the model parameters. Our simulations further reveal that population dynamics cannot be accurately described by the commonly-used surface growth models. Instead, one has to explicitly account for the emergent growth instabilities and demographic fluctuations. Overall, our work links together environmental conditions, colony morphology, and evolution. This link is essential for a rational design of concrete, biophysical perturbations to steer evolution in the desired direction.

## Introduction

Human life hinges on myriads of populations be it forests that produce timber or rhizobia that fix nitrogen [1]. To manage these populations, we need to understand how possible interventions could affect their ecology and evolution.

While there are many ecological studies that link environmental variables to growth patterns or spatial distribution of species, evolution is typically studied in more abstract settings that emphasize general principles rather than specific applications [2–4]. As a result, we lack practical knowledge of how to perturb the ecology of a species to slow down or accelerate its evolution.

The challenge of linking ecological and evolutionary dynamics can be most readily addressed in the context of microbial colonies [5–8]. Indeed, microbial growth is affected by only a handful of environmental variables, and nontrivial evolutionary changes occur already within the lifespan of a single colony [9–11]. Despite their simplicity, microbial colonies share many similarities with other cellular aggregates such as biofilms and tumors, which are of an immense practical importance [12, 13].

The ecological and evolutionary aspects of colony growth have, for the most part, been investigated separately. Ecological investigations focused on colony morphology and the effect of environmental variables such as nutrient concentration, which affects growth, and agar concentration, which affects motility [9, 14–16, 16–21]. Collectively, these studies established a phase diagram for colony morphology and proposed a minimal model that recapitulates many commonly observed spatial patterns. In contrast, evolutionary investigations focused almost exclusively on the simplest disk-like colonies. Moreover, theoretical models of disk-like colonies have typically imposed a certain type of dynamics (known as KPZ universality class or Eden model) that has not been derived from empirical data or from a mechanistic description of bacterial growth and motility [11, 22–26].

Here, we explore evolutionary dynamics in colonies of varied shapes and relate evolutionary processes to colony morphology and environmental variables. We adopt a classic model with a diffusible growth-limiting nutrient and nonlinear biomass diffusion [20]. We supplement this model with a stochastic component to account for demographic fluctuations and genetic drift. While the model reproduces a variety of colony shapes, our focus is on the two dominant morphologies: an approximately flat front and a rough front with irregularities due to growth instabilities. Using both analytical arguments and simulations, we elucidate how environmental variables affect colony roughness and genetic drift.

## Methods

Microbial colonies are convenient systems to study population dynamics. Microbial colonies are easy to grow and visualize, and numerous growth and dispersal strategies are available from a large pool of natural isolates and genetically engineered microbes. Given their ease-of-use, microbial colonies became model systems for other, more complicated, processes such as range expansions in ecology, biofilms in engineering, or tumor growth in medicine [12, 13, 23, 27, 28]. Microbial colonies are of course important in their own right, at least, because many empirical studies involve colony growth as an intermediate step in the experimental protocol. Not surprisingly, microbial colonies have been intensely studied by both theorists and experimentalists willing to understand the growth and form of perhaps the simplest of populations [17, 29–33].

Despite their apparent simplicity, microbial colonies could produce intricate spatial patterns that depend on both the species and the growth conditions [10, 16, 19, 34]. Some of the mostly commonly observed patterns include smooth and rough disks, concentric circles, scattered dots, and a whole zoo of branching morphologies that could even include chiral twisting [32, 35, 36]. Detailed experimental and theoretical studies, however, largely focused on the disk-like colonies and the transition between the disk-like and branched morphologies [20].

Historically, the motivation to simulate microbial growth was to determine the origin of the colony shape, in particular, the amount and type of roughness or undulations of the colony edge [17, 20, 29, 31]. The two primary mechanisms that were considered are demographic fluctuations and growth instabilities. Demographic noise creates small ripples along the colony edge, which can accumulate over time and result in a rough growth front. In contrast, growth instabilities produce finger-like protrusions that either result in well-developed branches or just in a rough growth front. The primary mechanism behind the growth instabilities is thought to be nutrient limitation [20], but mechanical origins of the instabilities have also been considered [37].

More recently, microbial colonies also became a prime system to study evolution and spatial population genetics [11, 23, 24, 38–43]. Most of the models that were used to interpret or explain experimental data, however, did not consider the true origin of demographic and evolutionary processes in microbial colonies. Instead, it was typically assumed the population dynamics can be described by a cellular automaton with simple growth rules (typically the Eden model [44]) or by a phenomenological theory that prescribed the motion of ancestral lineages to be either simple diffusion or super-diffusion driven by Kardar-Parisi-Zhang (KPZ) dynamics of the growing edge [22, 45, 46]. With this paper, we hope to make the first step in connecting the evolutionary processes in microbial colonies to their biophysical origins. To accomplish this goal, we chose to focus on demographic fluctuations and nutrient limitation as the primary drivers of colony shape. This is a reasonable choice because these two processes are so basic that they occur in nearly any microbial population. In addition, prior work has shown that models based on nutrient limitation can reproduce experimentally observed patterns remarkably well [14–16, 16–20].

### Nutrient-limited growth and self-activated dispersal

Before turning to a fully stochastic model, it is essential to understand the main ingredients of the deterministic dynamics, which are nutrient diffusion, nutrient consumption, growth, and motility. These processes are described by the following reaction-diffusion equations:

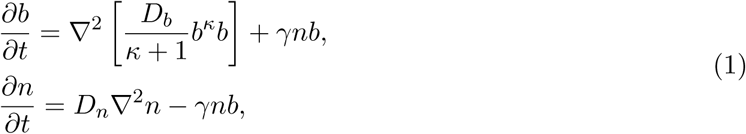

where *n* is the concentration of the growth-limiting nutrient, and *b* is the cellular biomass. Both quantities depend on time *t* and two spatial coordinates *x* and *y*. The rational for choosing this specific model is discussed in Refs. [20] and is also explained below.

The nutrient dynamics consists of diffusion with a diffusion constant *D*_*n*_ and consumption by cells at the rate of *γnb*. While more complex, e.g. Michaelis-Menten, rates of nutrient uptake can be considered, they are often not necessary to reproduce common colony shapes. Moreover, once nutrients are depleted, the dynamics are largely determined by the consumption rate at low nutrient levels, which is well approximated by *γnb*. All of the consumed nutrient results in the production of the biomass at the rate *γnb*. Thus, we neglected nutrients that are needed for basic metabolic maintenance and also set the conversion factor between nutrient concentration and biomass to unity. The latter is not an extra assumption because one can always measure nutrient concentration in the units of biomass that is produced from that nutrient.

In addition to the growth of the biomass, the model also describes its motility, parameterized by the motility constant *D*_*b*_ and the motility exponent *κ*. Standard diffusion is obtained for *κ* = 0, which is a good model of swimming motility. Cells grown on even moderately hard agar, however, cannot swim and need to move collectively either by pushing each other or by producing surfactant or lubricant [33, 34]. Both processes are cooperative and therefore become more efficient as the number of cells increases. This positive feedback is encoded by *κ*> 0. All simulations reported here are carried out for *κ* = 1 because neither we nor previous studies noticed significant qualitative differences when using other reasonable values of *κ*. Our theoretical derivations are however done for an arbitrary *κ* because this does not complicate the analysis.

Equations (1) and their modifications have been widely used to study microbial colonies because they exhibit growth instabilities, a classic mechanism of pattern formation that can produce different colony morphologies depending on the model parameters. A growth instability is a situation when the outward expansion of a flat front in unstable to even an infinitesimal perturbation. Many types of instabilities exist, but the type most relevant for microbial colonies is the so-called MullinsSekerka instability, which occurs when nutrient diffuse much faster than the biomass [20, 47**?**].

Nutrient diffusion destabilizes the growth because regions protruding forward gain greater access to nutrients and therefore tend to protrude even more while regions that caved in expand even slower because their access to nutrients is diminished. In contrast, biomass diffusion smooths out front undulations. When the biomass diffuses rapidly, it can overcome the variations in the growth rate and suppress the instability resulting in a relatively flat growth front. In the opposite limit, front develops bulges or fingers that are born due to demographic fluctuations or substrate inhomogeneities and amplified by the growth dynamics. Since the initial perturbations are random, the long-time dynamics is random as well. In fact, it could resemble the dynamics driven by demographic fluctuations on temporal and spatial scales much larger than those of the instabilities [48–50].

Growth instabilities in Eqs. (1) has been analyzed in Ref. [20] that demonstrated a transition from stable to unstable growth as 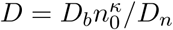decreased below a critical value of order unity. Similar transitions have also been obtained in related model [20, 33, 34].

## Simulations

Although Eqs. (1) provide an adequate description of colony growth, they do not account for demographic fluctuations and genetic drift. To probe these evolutionary forces, we sought a model that operates with several alleles (genotypes) and explicitly accounts for the stochastic nature of births and motility. Because stochastic simulations demand much greater computational power, we had to adopt a framework that balances biological realism with computational efficiency.

Given our interest in genetic drift, we modeled the dynamics of *K* neutral alleles, i.e. *K* distinct genotypes that have the same growth and motility. We denoted the biomass of these alleles as *b*_*k*_ and used a fixed time step *dt* and a square grid with a lattice spacing *dx* in both *x* and *y* dimensions.

Thus, nutrient and biomass concentrations were represented by matrices *n*(*i, j*) and *b*_*k*_(*i, j*) where *i* and *j* indexed the *x* and *y* dimensions of the spatial grid. Each simulation time step consisted of four updates: biomass migration, biomass growth, nutrient consumption, and nutrient diffusion; see Fig. 1a.

**Figure 1.**
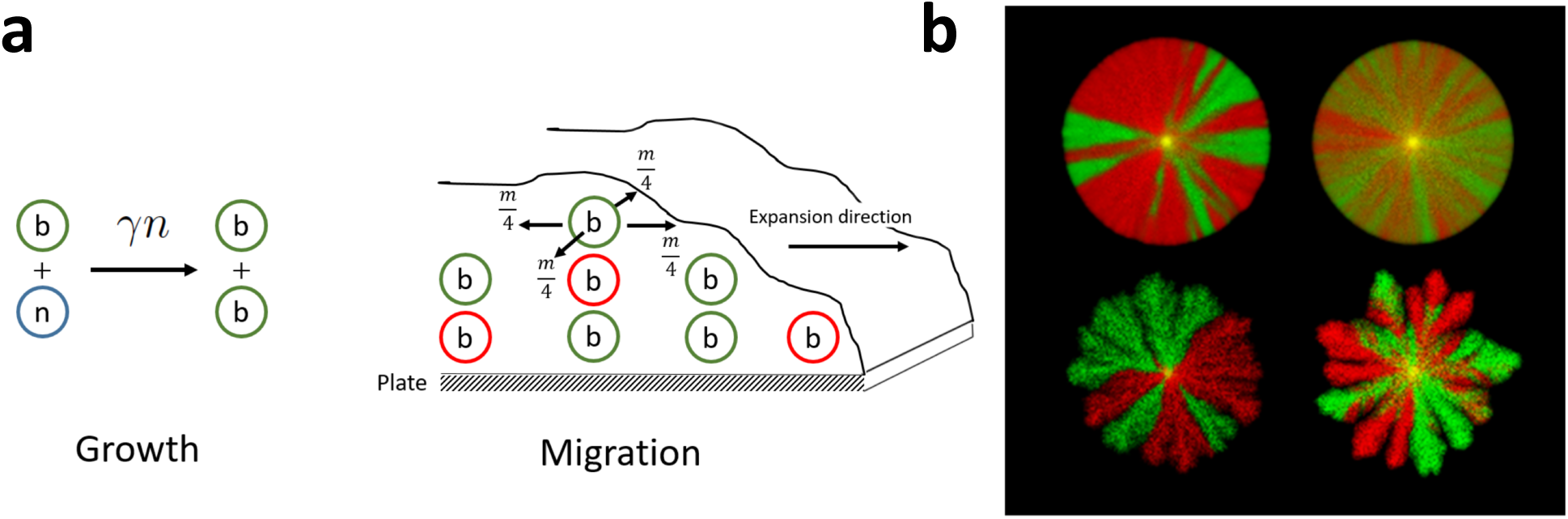
Nutrient limitation alters morphology and diversity of expanding colonies. **(a)** Colony expansion is due to growth and motility. Cells divide at a rate proportional to the nutrient concentration and migrate without any spatial bias; the migration rate depends on the biomass concentration and possibly on nutrient concentration. **(b)** Our model can reproduce many of the typically observed colony morphologies including circular disks and colonies with finger-like protrusions or branches. Note that similar morphologies could have very different rates of genetic drift. This is evident from the number, crispness, and motility of boundaries between two different genotypes labeled with red and green colors. Starting from well-mixed initial conditions (yellow centers), the monochromatic domains (sectors) emerge because one of the genotypes becomes extinct locally due to the vagaries of reproduction and migration at the front.

Biomass growth was controlled by the division rate *g*(*i, j*) that could in principle have an arbitrary dependence on biomass and nutrient concentrations. Because divisions are discrete events, we needed to convert cellular biomass into the number of cells. This conversion was accomplished by dividing the total biomass *b* by the biomass of a single cell Δ*b*. Given the division rate *g*(*i, j*) and the total biomass *b*(*i, j*), the number of divisions at that site is binomially distributed with *g* being the success rate and *b*/Δ*b* being the number of trials. We then distributed these divisions among the *K* alleles in proportion to their relative abundance using a multinomial distribution. Because nutrient consumption and biomass growth are interlinked in Eqs. (1), we reduced *n*(*i, j*) by the biomass of a single cell, Δ*b*, for each cell division. This was a convenient and efficient implementation even though it introduced unnecessary stochasticity in the nutrient dynamics. For all simulations presented in this paper, we used *g*(*i, j*) = *γn*(*i, j*) following Eqs. (1).

Biomass motility was modeled by migration of cells between the grid points. The migration rate (per cell) out of site (*i, j*) was set to

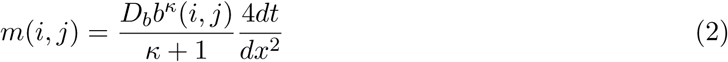

in order to reduce to Eqs. (1) in the continuum limit. The total number of migration events and their distribution among the *K* alleles were carried out just as for the biomass. Each migrant moved to one of the nearest neighbors with equal probability. We also had to supplement Eq. (2) with an additional rule to account for the reduced motility behind the growing edge.

Typically, in the bulk of the colony, the cells are densely packed, and the nutrient concentration is very low [5, 23]. As the results, cells become nonmotile or even enter a dormant state. This cessation of motility is not relevant to the overall colony morphology, which is primarily determined by the cell motion at the colony edge. Therefore, prior work leading up to Eq. (1) did not consider changes in motility within the bulk of the colony. The cessation of motility is however important for an accurate description of sector or domain boundaries that form between alleles as the colony grows [11]. Otherwise, the sector boundaries would blur over time and eventually all parts of the colony would intermix. We implemented motility cessation by setting *m*(*i, j*) to zero once *n*(*i, j*) fell below a critical value of *n*_*c*_ = 0.01.

We also briefly explored motility rates that were proportional to the product of nutrient and biomass concentration. This choice has been used previously to model certain bacteria and reflects the dependence of motility on the number of actively growing cells [51]. We found that cell-free grooves that develop under low nutrient conditions occur only when biomass motility increases with both *b* and *n*. Otherwise, the behavior of the models was similar, so we focused on the simpler and more widely-used model of nutrient-independent biomass diffusion.

Finally, the diffusion of the nutrient was modeled deterministically by solving the diffusion equation using the Crank-Nicolson method [52]. The use of this implicit method allowed us to take a relatively large *dt*, which would violate the stability condition in an explicit method, and thus dramatically speed up the simulations.

We show representative simulations of whole-colony growth in Fig. 1b. Genetic drift is illustrated with *K* = 2 alleles that are labeled in two different colors. It is easy to see that by varying simulation parameters one can obtain a wide range of morphologies and genetic behaviors. The colony edges range from very smooth boundaries to non-space filling tendrils while sector boundaries between different genotypes range from very sharp to very fluid.

Although whole-colony simulations are pleasant to the eye, they are inefficient because the colony edge occupies only a small fraction of the simulated spatial grid. We overcame this problem by focusing on linear inoculations, which are typically done with a razor blade or a similar object. Thus, we considered a front that is flat initially and modeled its motion in a rectangular simulation box. Since the growth of the colony is limited to a narrow region near its edge, we avoided the need to simulate an entire colony by moving the simulation box with the growing front. This “treadmilling” was accomplished by shifting the biomass and nutrient profiles once the biomass occupied close to half of the simulation box. The biomass shifted outside of the simulation box was processed and recorded for further analysis. The biomass shifted into the simulation box was set to zero, and the nutrient shifted into the box was set to *n*_0_, the initial nutrient concentration in the Petri dish. For simplicity, we used periodic boundary conditions in the direction transverse to the colony growth. The length of the box in this transverse direction is denoted as *L* and represents the latteral spatial extent of the expansion front.

Except when indicated otherwise, we used *D*_*n*_ = 0.1, *γ* = 0.01, *L* = 300, Δ*b* = 10, *dt* = min{1, *dx*^2^/(16*D*_*b*_*n*_0_)}, *κ* = 1, *dx* = 0.25, and averaged over 20 replicates.

### Dimensional analysis

In the continuum limit, our simulations can be described by the following set of stochastic partial differential equations:

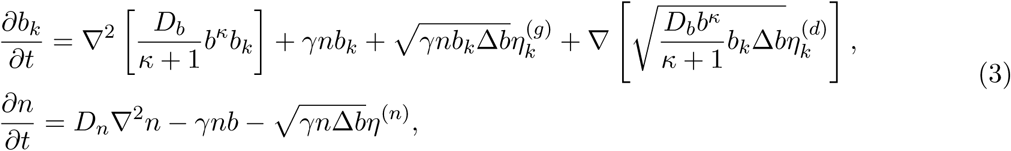

which, except for the noise terms, are identical to Eqs. (1). Note that we did not include the cessation of migration into these equations. The reason for this is that all of the quantities of interest to us here can be derived from *b*_*k*_ measured only at the growing edge and thus do not depend on the dynamics in the colony bulk. By not accounting for the dynamics behind the front, we made the following analysis much more straightforward.

The new stochastic terms reflect fluctuations due to randomness in the growth (labeled with ·^(*g*)^) and dispersal (labeled with ·^(*d*)^). The former accounts for both the randomness in the division times (demographic fluctuations) and the randomness in the type of cell that is chosen to divide (genetic drift). The latter accounts for the randomness in the choice of whether to migrate or stay at the same location and the randomness in the choice of the migration direction. Because migration does not alter the number of cells, the integral of the dispersal noise over space must be zero, which can be easily demonstrated by integrating by parts.

Each stochastic term is a product of its strength and a normalized Gaussian white noise (labeled by various *η*), which are zero mean and delta-correlated. For example, the statistical properties of the growth noises are as follows

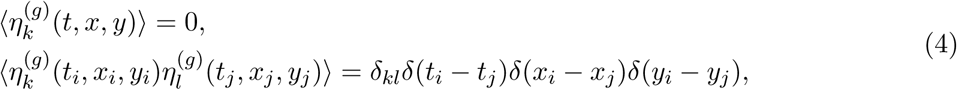

where the angular brackets denote averaging over the realizations of the noise. The dispersal noises have identical statistical properties and are independent of the growth noises. The noise in the nutrient equation is discussed below and is derived from 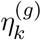.

The strength of the noise terms can be derived from the microscopic description of our simulations and is consistent with the prior analysis of similar models [53–57]. In particular, the strength of the growth noise is given by the rate of births (there are no deaths in our simulations), and the strength of the dispersal noise is given by the diffusion coefficient times the population density. The only unusual term in these equations is Δ*b*, the biomass of a single cell. This is just a normalization factor that accounts for the fact that *b* models the biomass instead of the number of cells.

The noise in the nutrient equation follows directly from the biomass equation because the only source of stochasticity is the consumption of nutrients during cell division. In fact, *η*^(*n*)^ is definedsuch that 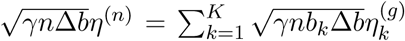 With this definition, it is easy to show that *η* is a zero-mean, delta-correlated Gaussian noise, just like 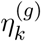 in Eqs. (4). We note that the noise in the nutrient equation is the consequence of our choice to consume nutrients in discrete amounts in our simulations. A more realistic, continuous consumption would produce much weaker noise. However, given our focus on the dynamics of the cells rather than the nutrients, this is a negligible artifact of our simulation scheme.

The dynamical equations (3) need appropriate boundary and initial conditions. Initial amount of the biomass is assumed to be small, and it does not affect the medium to long term dynamics considered here. In contrast, the initial nutrient concentration *n*_0_ plays a major role because it controls the rate of colony growth and the population density. The boundary conditions depend on the geometry of the expansion and directly follow from the description of our simulations. For linear expansions, the finite spatial extent of the simulations introduces another model parameter: *L*, which is the spatial extent of the front perpendicular to the expansion direction. This length scale is relevant for some quantities such as the time to extinction of all but one allele.

As stated, the mathematical description by Eqs. (3) contains seven parameters: *D*_*n*_, *D*_*b*_, *κ, γ*, Δ*b, n*_0_, *L*. This number of parameters overstates the true complexity of the model because solutions typically depend on a smaller number of certain parameter combinations. These combinations can be uncovered by the standard procedure known as dimensional analysis or scaling.

The idea behind dimensional analysis is that problems at different scales are related. For example, one can determine the weight of the Eiffel Tower by weighing its exact copy manufactured one thousand times smaller. The weight of the actual tower would be the weight of the copy times one thousand cubed because the weight of a three dimensional object scales with the cube of its linear size.

Similar procedure could be applied to dynamical equations like Eqs. (3) by introducing new scaled variables:

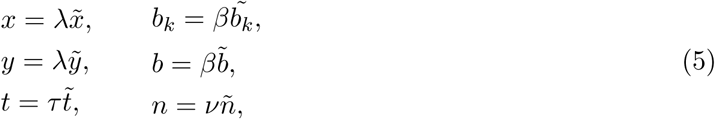

substituting them into Eqs. (3) and then setting to unity as many coefficients as possible in front of various terms. This calculation yields the values of the scaling constants λ, *τ, β, v* and the transformed equations. We find that

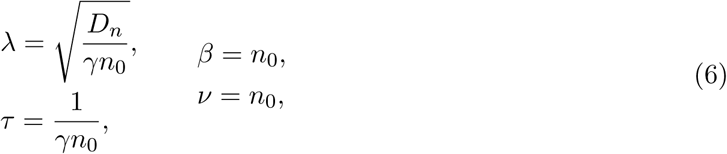

and the rescaled equations take the following form

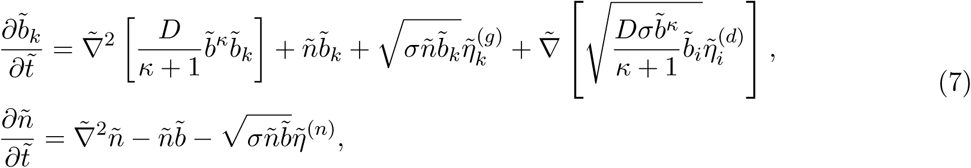

where 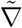denotes derivatives with respect to 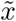 and 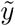, and the tildes above the noise variables *η* indicate that they are delta-correlated in the rescaled variables. Also, note that, the initial nutrient concentration equals one, and the system size (transverse extent of the front) equals *L*/λ after rescaling.

The two new dimensionless parameters in Eqs. (7) are given by

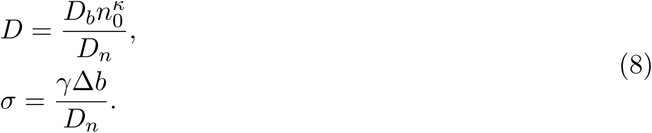

These are the primary factors that control qualitative aspects of colony growth and evolution. In particular, *D* controls the onset of growth instabilities and *σ* the strength of demographic fluctuations and genetic drift.

From our dimensional analysis, it is clear that the behavior of the colony is controlled only by: *D, σ*, and, for some quantities, by *L*/λ. The last parameter, *L*/λ, becomes important only for long-enough simulations, and thus many quantities are controlled by only two parameters. This is a significant simplification compared to the initial model (Eqs. (3)).

Once the solution for the rescaled equations (7) is obtained one can determine the solution of the original problem by reversing the rescaling transformation. Often times, however, one is interested not in the values of *b*_*i*_ and *n*, but in derived quantities such as the extinction time or sector width. Such quantities can be directly obtained from the rules of spatial and temporal scaling specified by Eq. (6). For example, the time to fixation of one of the alleles reads

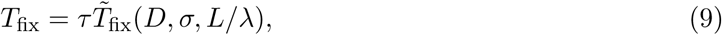

and the average size of sectors, *l*_*s*_, at some time *t* is given by

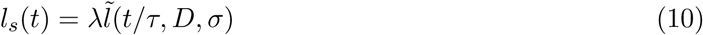

assuming that *l* ≪ *L*, so that there is no dependence on the system size.

### Morphology

The morphologies that arouse from linear inoculations are well-characterized by the position of the colony edge. We defined the edge as the first point in the direction of the expansion (*y−*axis) where the biomass fell below a predetermined threshold *b*_*c*_ = Δ*b*. For historical reasons [46], the edge is often referred to as height, and its *y−*coordinate is denoted as *h*. The exact location of the edge of course varies along the front (*x−*axis) and changes with time, so the shape of the colony is encoded in *h*(*t, x*).

From *h*(*t, x*), we can derive a few quantities that characterize the motion and shape of the colony. These are obtained by averaging over spatial variations along the front (denoted with an overbar) and, if needed, by averaging over the realizations (denoted with angular brackets). The spatial averaging can be done globally across the whole system (*x* varies from 0 to *L*), but it is often more convenient to perform a local average over a smaller region, say, from *x*_0_ to *x*_0_ + *l*. For local averages, we typically also average the result over *x*_0_, which can be done easily because of the periodic boundary conditions. In the following, we provide expressions for local averages; the global averages can be obtained by replacing *l* by *L*.

The most basic description of the colony motion is the average position of its front:

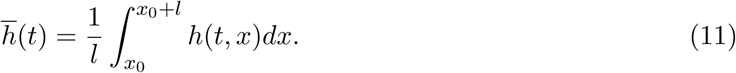

The time derivative of the average height gives the velocity of the front:

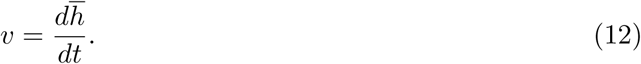

The degree of front undulations, which is typically called roughness [46], is captured by the variance of *h*(*t, x*):

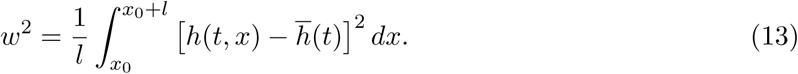

The above quantities are specific to each realization. In this paper, we only use their average over the realizations: ⟨*v*⟩ and ⟨*w*^2^⟩. We refer to them simply as *v* and *w*^2^ not to complicate the notation unnecessarily.

The estimate of the velocity does not depend on the size of the averaging window *l* (at least to the first order), but the roughness *w* typically increases with *l* unless there are strong mechanisms to suppress front undulations. The origin of this dependence could be multifactorial [46]. In rough colonies dominated by demographic fluctuations, the front performs a type of a random walk in *x*, so the increase of *w* with *l* is analogous to the increase of the root-mean-square displacement with time. Growth instabilities is another common factor. On small spatial scales, the front could be approximately flat, but located at an angle to the *x* axis. Thus the total width of the front grows very rapidly (approximately linearly) with *l* until *l* exceeds the instability length scale. The roughness then grows more slowly due to the undulations of the front on large spatial scales that could, for example, span several finger-like protrusions.

In many physical systems, the dependence of *w* on *l* and *t* is well-described by a power law [46]. For the simplest case of non-anomalous roughness, *w* grows as *t*^*β*^ at early times, but then saturates to a value that increases with the window size as *w* ~ *l*^*α*^. These scaling laws have been fitted to bacterial colonies previously even though the range of possible power law behavior is small. For comparison with previous work, we estimated the values of these exponents using a linear fit on a log-log plot.

### Diversity

Without mutations, diversity is lost due to genetic drift. Genetic drift is a term for fluctuations in the relative abundance of alleles, which can push some genotypes into extinction [4, 11]. Thus, the rate of diversity loss is a measure of genetic drift, a potent evolutionary force.

In spatially extended populations, genetic diversity is a function of both time and space. Because genetic drift is local, different spatial regions experience different fluctuations at least until migration of cells reduces these differences. These dynamics often manifest in the formation of sectors, which is a commonly-used method to visualize evolutionary dynamics in microbial colonies [11, 23]. Initially, the colony is a mixture of two or more equally-fit strains each expressing a unique fluorescent protein. As the colony growth proceeds, genetic drift drives the fixation of different strains locally and produces genetically homogeneous regions (sectors) separated by sector boundaries; see Fig. 1b. Once sectors are formed the subsequent dynamics is primarily driven by the motion of the sector boundaries, which is in turn controlled by the shape of the front. Indeed, the outward growth of the colony, makes sector boundaries move “downhill”, i.e. in the direction opposite to the spatial gradient of *h*(*t, x*) [22, 58–60].

The early stage of genetic demixing can be characterized by the average local heterozygosity, which is the probability to sample two different genotypes at a given location [11]. Mathematically, it can be expressed as one minus the probability to sample two identical genotypes:

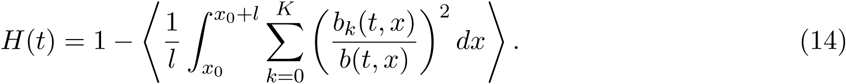

If alleles are equally-abundant initially, then *H*(0) is close to 1*−*1*/K*. Once sectors are formed, *H*(*t*) is small and approximately given by the ratio of the sector boundary thickness to sector width [11].

The sector boundary thickness does not change with time because it is determined by the balance of genetic drift and migration [53]. Thus, the continuing decrease of *H* is driven by the increase in the sector size due to the loss of small sectors when their two boundaries merge. The rate of these mergers depends on how sector boundaries move. A typical assumption, which has some theoretical and experimental support [22, 23, 30, 42, 61], is that the boundary moves as a diffusive or superdiffusive random walk with the root-mean-square displacement growing as *t*^*ζ*^, where *ζ* = *β/α* for models that satisfy the corresponding power-law scaling for *w*. Under this assumption, *H*(*t*) ~*t*^−*ζ*^ [11, 22, 42].

To probe the wandering dynamics of sector boundaries, we examined the stochastic changes of allele frequencies in simulations with pre-formed sectors. At *t* = 0, the front was divided into *K* = 20 equally-sized spatial domains. Each spatial domain was filled with a unique allele. Thus, the initial frequency of all genotypes was *f*_*k*_ = 1*/K*. These frequencies changed due to the motion of the two sector boundaries surrounding each allele. Therefore, up to a numerical factor, we can interpret the standard deviation of *f*_*k*_(*t*) *− f*_*k*_(0), averaged over *k* and different simulations, as a proxy for boundary wandering. Note that to convert from allele frequencies to boundary displacement, we multiplied the standard deviation of *f*_*k*_(*t*) *− f*_*k*_(0) by the width of the expansion front *L*.

The connection between boundary motion and *f*_*k*_(*t*) is strictly valid only at early times when the boundaries move independently from each other because their spatial separation exceed the correlation length associated with front roughness. When boundary separation becomes small, e.g. right before the extinction of the allele, the motion of the boundaries become correlated and therefore not directly related to the changes in *f*_*k*_. In our simulations, the contribution of sectors approaching extinction was small, and we do not expect it to significantly affect the qualitative nature of boundary wandering. Therefore, we chose the easy-to-quantify (and therefore robust) metric of root-mean-square changes in *f*_*k*_(*t*) as a measure of boundary wandering instead of trying to extract the spatial positions of each boundary, which would necessitate additional assumptions in order to convert the spatial distribution of genotypes into the positions of sector boundaries.

## Results

The minimal mechanistic model developed above allowed us to ask how various environmental and cellular properties affect population dynamics in microbial colonies. Dimensional analysis suggests that both genetic drift and colony shapes are controlled by only two quantities: *D* and *σ*. The first quantity, 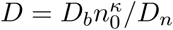 depends on biomass diffusion *D*_*b*_ and initial nutrient availability *n*_0_, which can be easily controlled experimentally by varying the agar and glucose concentrations respectively. In addition, prior work predicted that the transition from stable to unstable growth occurs as *D* is increased beyond a critical value of order unity [20]. In contrast, the second quantity *σ* = *γ*Δ*b/D*_*n*_ depends on parameters, which are much harder to vary experimentally. Furthermore, *σ* just determines the strength of demographic fluctuations, and there is nothing to indicate that the magnitude of the noise can lead to a qualitative change in the behavior. Therefore, we focused on exploring the effects of varying *D* by changing the nutrient concentration or bacterial motility.

The primary goal of numerical simulations was to answer two related questions. First, we wanted to confirm whether the predictions of dimensional analysis hold that is to verify that simulations with the same values of *D* exhibit the same behavior up to the scaling transformation (Eq. (6)). Second, we wanted to determine how the morphological transition from rough to smooth colony edge affects genetic drift and genetic diversity.

Our approach to answer both of these questions is illustrated in Fig. 2, which shows that the *n*_0_-*D*_*b*_ plane is divided into two regions: unstable, strongly undulating fronts at low *D* and stable, nearly flat fronts at high *D*. Given the computational constraints of our simulations, we found model parameters that allowed us to accurately simulate population dynamics for a very low value of *D*. We then increased the value of *D* either by increasing the biomass motility (orange) or by increasing the nutrient concentration (blue). Although, we explored a wider range of *D*, our detailed analyses are focused on the three pairs of simulations each with the same *D* = 0.05, 0.5, or 1, but different *D*_*b*_ and *n*_0_. Note that all of the simulations have the same *σ* because we kept *D*_*n*_, Δ*b*, and *γ* fixed.

**Figure 2.**
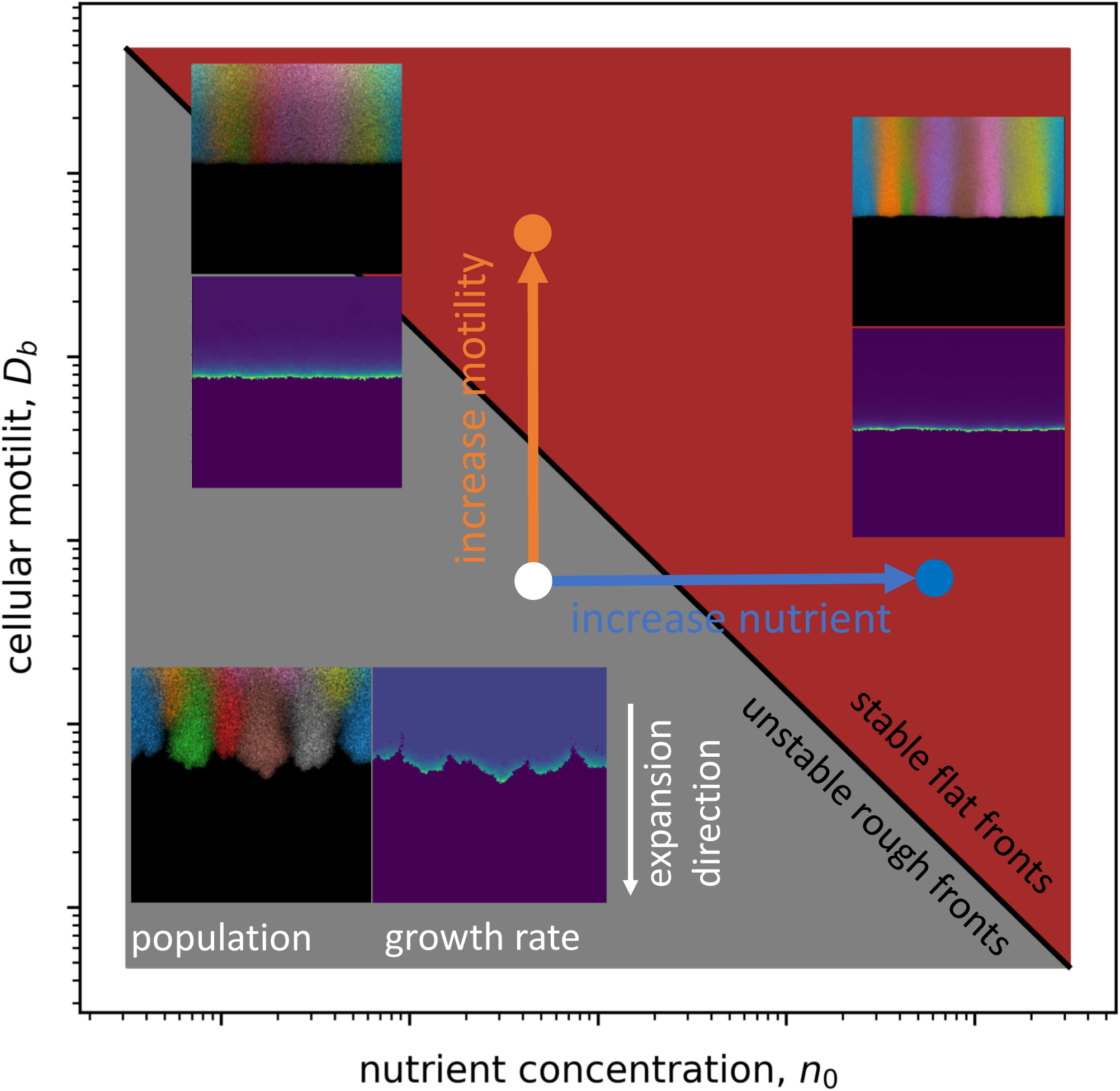
Colonies transition from rough to smooth as nutrient concentration or cellular motility are increased. This schematic illustration shows that we investigate how the morphology and genetic diversity of the colonies change by varying either *n*_0_ or *D*_*b*_. One set of insets shows the shape and genetic composition of the front. The other set of insets shows the spatial variation of the growth rate. Note that the two flat front have different depth of the growing layer and different sizes of sectors.

Figure 2 shows dramatic changes in the morphology and genetic diversity as *D* increased in our simulations. Furthermore, this figure highlights major differences between simulations with the same value of *D*. Specifically, the number of sectors is much lower and sector boundaries are much more diffuse in the simulations with higher motility compared to the simulations with more nutrient. The differences between simulations are also evident in biophysical metric such as the spatial distribution of actively growing cells, which, of course, influences genetic processes within the colony [5].

If our dimensional analysis is valid, these differences between simulations with the same *D* should only be a matter of spatial and temporal scales specified by Eqs. (6). In the following, we directly test this prediction for practically relevant morphological and demographic metrics by comparing population dynamics in original and rescaled variables.

### Expansion velocity

Expansion velocity is the most basic and practically important metric of colony growth. We found that increasing either biomass motility or nutrient concentration sped up the colonization rate (Fig. 3a). The amount of the speedup was higher when nutrient concentration was increased than when biomass motility was increased The greater effect of *n*_0_ is expected from the dimensional analysis because 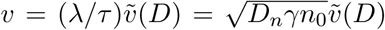 Indeed, increasin *D*_*b*_ influences *v* only through 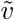 while increasing *n*_0_ increased both 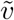 and the scaling prefactor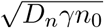 The validity of the dimensional analysis was further confirmed by rescaling both time and space according to Eqs. (6) and observing that the expansion dynamics in 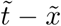 coordinates is exactly the same for simulations with the same values of *D* (Fig. 3b).

**Figure 3.**
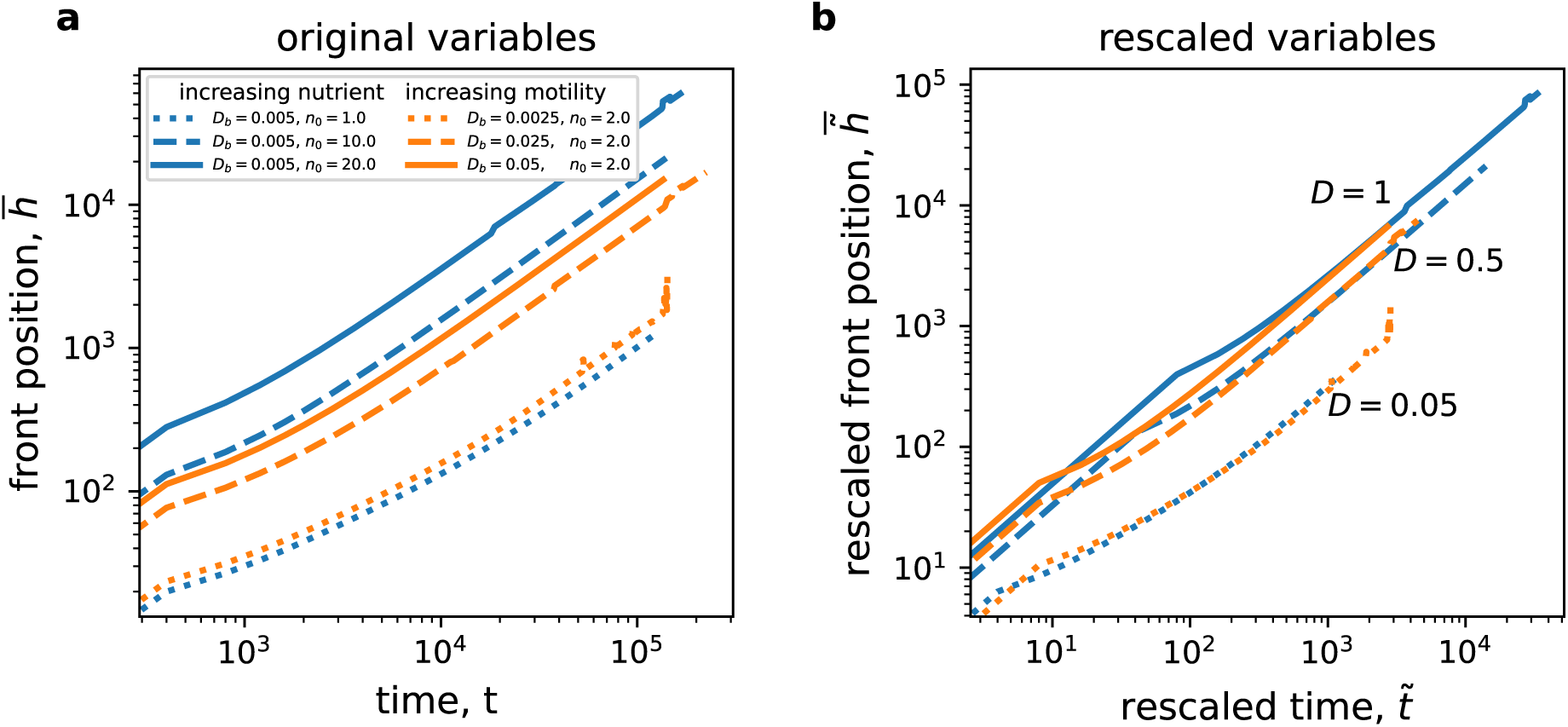
Expansion velocities agree with dimensional analysis. After an initial transient, the expansion proceeds with a constant velocity, i.e. 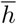 is linear in *t*. Data with the same value for *D* are shown with the same linestyles (dotted, dashed, and solid), and the color indicates whether we changed the nutrient levels (blue) or the motility (orange). Original variables, without rescaling, are used in **(a)**. It is clear that both *n*_0_ and *D*_*b*_ increase the expansion rate, although the nutrient level has a greater effect. Panel **(b)** demonstrates that *D* is the only factor controlling the expansion dynamics after rescaling according to Eqs. (6).

### Colony roughness

We then examined the effect of varying model parameters on colony shapes by computing the roughness of the expansion front; see Eq. (13). Since linear expansions were started with flat fronts, the roughness was essentially zero initially, but it increased dramatically over time. This increase consisted of two qualitatively different regimes. At early times, the roughness grew rapidly because of the growth instabilities that created characteristic undulations or even protrusions along the front. The roughness then grew more slowly once the sizes of undulations stabilized, and the front shape changed primarily via the birth and death of protrusions.

Given this difference between early and late time dynamics, a simple power law scaling (see Methods) cannot describe roughness in microbial colonies. The early time dynamics occurred over such a short time that a power-law fit may not be very meaningful. Furthermore, there were substantial differences in the apparent slope (*β*) on the log-log plot among simulations with different parameter values. A typical value of this slope was around 0.7 *−* 1.1, which is inconsistent with the predictions of the KPZ and Edwards-Wilkinson (EW) universality classes, but close to the prediction of the qKPZ universality class and previous theoretical and numerical studies [17, 45, 46, 49]. The late-time dynamics had a significantly lower slope *β ≈* 0.14, which was also inconsistent with the predictions of any of the classic models.

The dependence of roughness on *D* was much simpler. For any *t*, we found that roughness decreased with *D*, i.e. the fronts became flatter and the magnitude of growth instabilities decreased (Fig. 4a). The reduction of front undulations was more dramatic upon increasing the nutrient concentration compared to increasing the biomass motility. This is again consistent with the dimensional analysis, which predicts that 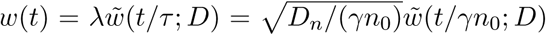 i.e. higher nutrient concentration reduces *w* through the scaling prefactor, *D*, and by decreasing the rescaled observation time while *D*_*b*_ only affects *D*. We also confirmed the predictions of the dimensional analysis quantitatively by checking for the collapse of roughness plots for the same values of *D* in the rescaled coordinates (Fig. 4b).

**Figure 4.**
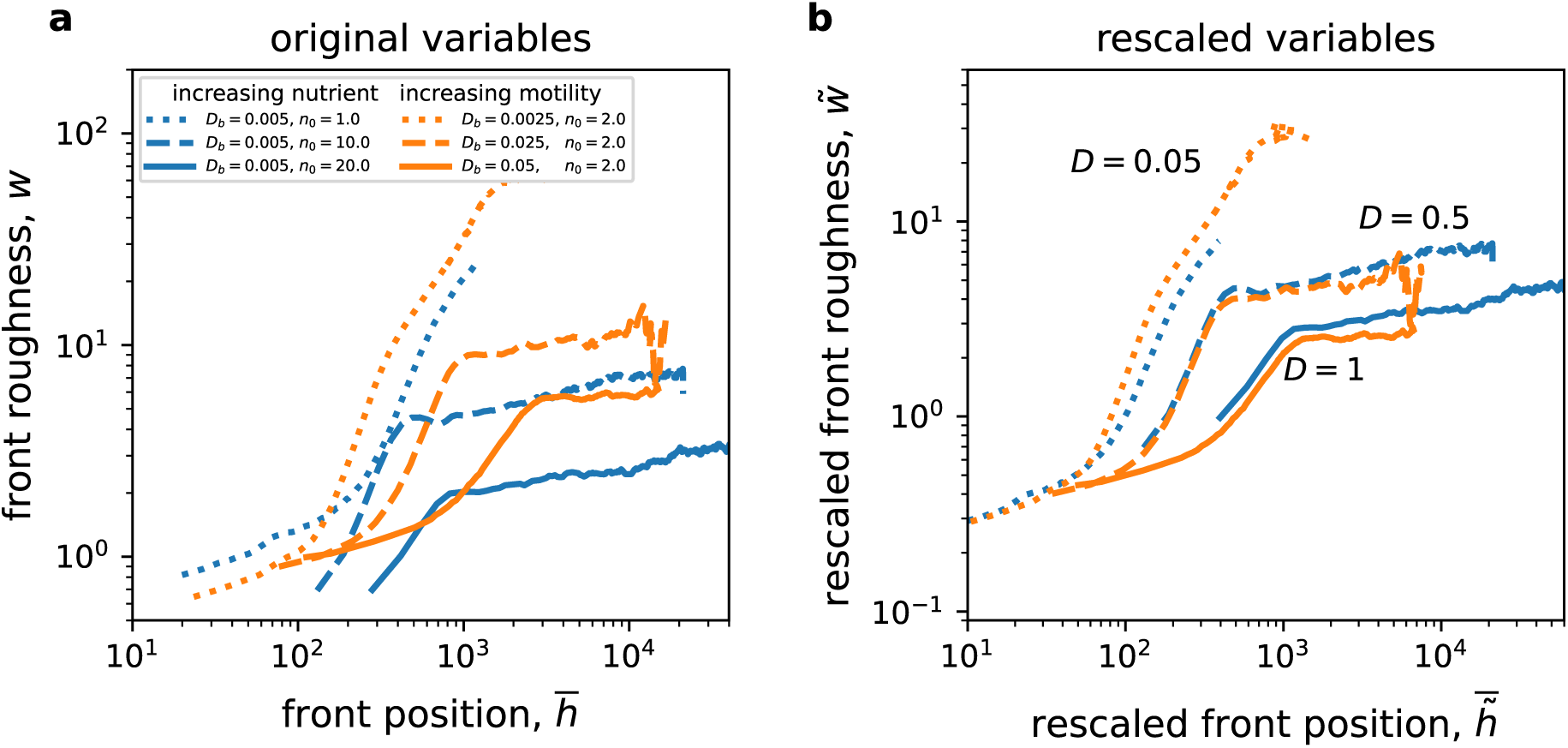
Front roughness decreases with nutrient concentration and cellular motility. Front roughness, *w*, measures the width of front undulations; see Eq. (13). As the expansion proceeds, the roughness first increases rapidly, as the growth instabilities emerge, and then more slowly. We show this increase as a function of the front position 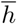 instead of time because the former is easier to obtain from experimental images and allows direct comparison between colonies of similar size. As in Fig. 3, panel **(a)** uses original and panel **(b)** rescaled variables. The collapse of the curves onto each other in panel **(b)** validates our dimensional analysis. Both *n*_0_ and *D*_*b*_ reduce roughness, but nutrient concentration has a stronger effect.

The roughness collapse was less precise then the collapse of front positions in Fig. 3 presumably because of the discreteness of space and time, which is present in our simulations, but not in the theory or experiments. Given that front shape changes on small length scales, it is indeed reasonable to expect that discreteness has a greater effect on *w* compared to *h*. The quality of the collapse is also influenced by the degree of averaging. Velocity, being a deterministic quantity, can be computed accurately from just a handful of simulations. In contrast, roughness is much more variable, so the estimate of the average roughness has a larger statistical error given the same number of simulations.

### Diversity loss

The decrease of growth instabilities with higher values of *D* also had profound effects on the genetic diversity in the simulated colonies.

We first explored genetic demixing that leads to the formation of sectors by starting our simulations with just two genotypes that were intermixed along the front. This mixed state became more segregated over time, which is evident from the decay of local heterozygosity *H*(*t*) in Fig. 5a. The decay was more rapid at low values of *D* suggesting that growth instabilities promote genetic demixing. Indeed, only portions of the front that significantly protrude forward can contribute to future generation in irregular fronts. This naturally reduces the effective population size and therefore increases the strength of genetic drift.

**Figure 5.**
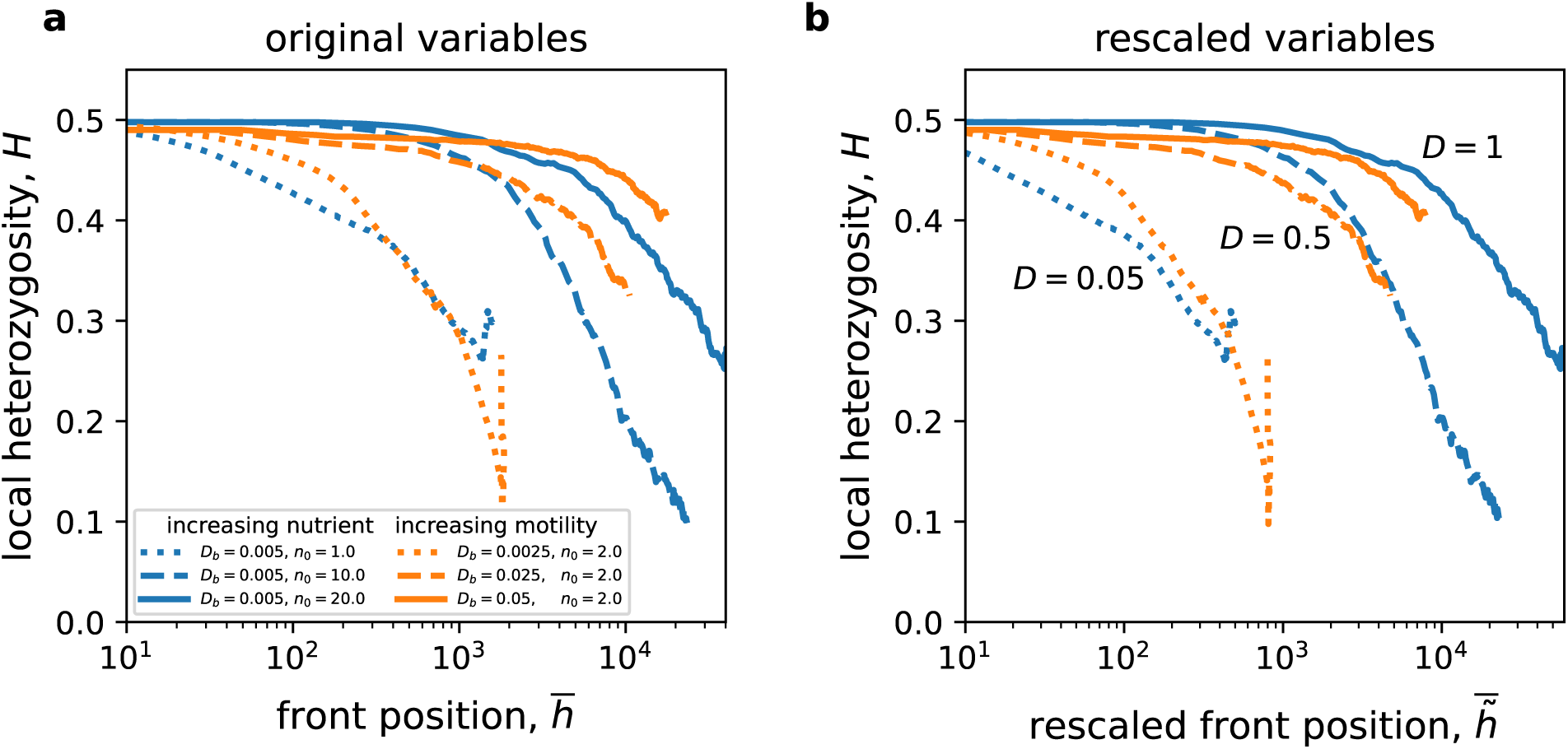
Rough fronts demix more rapidly. Genetic diversity is quantified by the average local heterozygosity (see Eq. (14)), which is the probability to sample two different genotypes at any spatial locaton. The simulations were started with well-mixed populations with two alleles. Over time, genetic drift promotes sectoring via local extinction of the genotypes. This manifests in the decline of the heterozygosity. Rougher fronts with lower value of *D* lost diversity more rapidly. The separate role of *n*_0_ and *D*_*b*_ is less clear than in the previous figures, but the data strongly suggests that populations with higher motility preserved the diversity better than populations with higher nutrient concentration. There is also modest evidence that rescaling improves the collapse of the curves.

Although there was a clear dependence of *H* on *D*, there were only minor differences between simulations that achieved a given value of *D* by varying either *D*_*b*_ or *n*_0_. Thus, it was difficult to determine whether rescaling improved the collapse of *H*(*t*) plots for the simulations with the same *D*.

We then explored genetic drift on longer times scales, after the formation of sectors. In this regime, the changes in allele frequencies are controlled primarily by the motion of sector boundaries. Therefore, we started the simulations with preformed, equally-sized sectors each with one of *K* = 20 different alleles and examined the wandering of sector boundaries by tracking changes in allele frequencies. The results of these simulations are shown in Fig. 6. As with the local heterozygosity, we found that boundary wandering and therefore genetic drift were stronger for rougher fronts (lower values of *D*) and that the differences between simulations with the same value of *D* were too small to evaluate the effects of the rescaling transformations.

**Figure 6.**
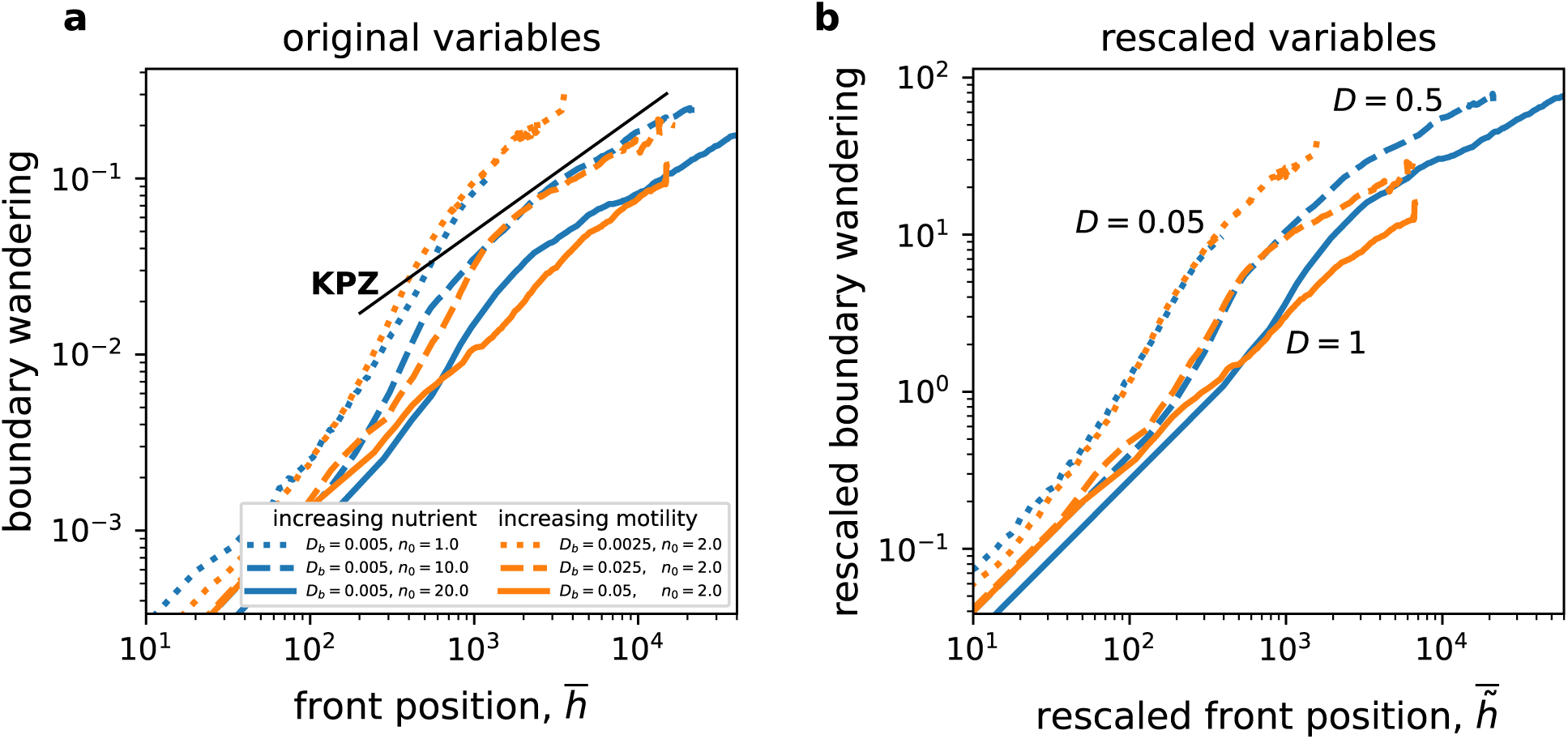
Boundary wandering shows dynamics expected for the KPZ universality class.. The motion of sector boundaries is quantified by the root-mean-square of the changes in allele frequencies. Similar to roughness, there appears to be two distinct temporal phases, although quantitative differences between the phases are less striking. The magnitude of boundary wandering decreases with *D*, and both *n*_0_ and *D*_*b*_ seem to have a similar effect. Since there is little difference between the curves with the same *D*, it is not possible to say whether rescaling in panel **(b)** results in a better collapse compared to original variables in panel **(a)**. The black line indicates a slope of 2/3, which is the expected exponent for the KPZ universality class.

Similar to roughness, boundary wandering consisted of two distinct phases. However, the quantitative differences between the early and late stage dynamics were somewhat smaller. In the late-time phase, the typical displacement of sector boundaries increase as an approximate power law of time with an exponent that did not change with *D*. The value of this exponent was close to 2/3, which is predicted by the standard models of surface growth [17, 25]. The boundaries moved more rapidly during the transient.

Thus, the motion of sector boundaries in our model was very similar to boundary motion in models that do not explicitly model nutrient depletion and thus do not exhibit growth instabilities. Given that the standard models did not describe the increase of colony roughness well (at least not in the late-time regime), one should be caution about assuming that the KPZ equation can describe all aspects of colony growth even if it provides a good description of boundary motion.

## Discussion and Conclusions

Microbial biofilms, cancer tumors, and other cellular aggregates are often difficult to control because they evolve rapidly in response to external perturbations [13, 62]. Genetic diversity is thought to be a major component of this high potential for adaptation, and spatially structure often allows for much higher levels of genetic diversity compared to well-mixed conditions [11]. Therefore, it is essential to understand how genetic diversity depends on the biophysical aspects of the environment and how it relates to the shape of a cellular aggregate. We explored these questions using a minimal model of a microbial colony, which, apart from growth, includes diffusion of a growth-limiting nutrient and cooperative motility on a hard surface. This model captures many biophysical aspects of colony growth and recapitulates several of the commonly-observed colony morphologies [16, 20].

Despite its overall simplicity, the model contained a large number of parameters, many of which could differ among species or environmental conditions. Dimensional analysis greatly reduced this complexity and determined that any quantity of interest is a product of an easy-to-compute scaling factor and an unknown function of just two dimensionless parameters *D* and *σ*. The former controlled the onset of growth instabilities and the latter represented the strength of demographic fluctuations. We confirmed the functional forms predicted by the dimensional analysis using simulations with different parameter values that nevertheless resulted in identical values of *D*. Such simulations exhibited identical behavior once we rescaled spatial and temporal variables by the appropriate scaling factors.

Out of the many model parameters that can in principle be modified, we focused on the nutrient concentration *n*_0_ and biomass motility *D*_*b*_ that can be easily modified experimentally by changing the glucose and agar concentrations respectively. Both of these parameters affect *D* equally since *D ~ D*_*b*_*n*_0_ (for *κ* = 1). Nevertheless, we found that nutrient concentration had a much greater affect on the expansion velocity and colony roughness because the relevant scaling factors depend on *n*_0_, but not *D*_*b*_. For genetic diversity, the contribution of rescaling factors was small, and the differences among the simulations were largely explained by the value of *D*, i.e. *n*_0_ and *D*_*b*_ contributed about equally.

Overall, *D* emerged as the main parameter affecting both diversity and morphology. Indeed, *D* controls the onset of growth instabilities, so near the transition from nearly flat to rough fronts, the size and pace of growth instabilities should depend on *D* very strongly. As a result, there is a strong correlation between genetic diversity and colony shape: Rough fronts typically have fewer and larger sectors compared to flat fronts. This, however, is not an absolute rule. Figures 4 and 6 show an example of two conditions (solid orange and dashed blue) that have the same roughness, but very different extent of boundary wandering. Thus, in general, one cannot quantify genetic processes from colony morphology alone. This is especially true for flat fronts that can appear very similar morphologically but nevertheless experience very different rates of genetic drift (Fig. 2).

Interpretation of colony shapes and genetic diversity is further complicated by significant differences between the early times when growth instabilities emerge and later times when growth instabilities at different spatial locations compete with each other. For both front roughness and boundary wandering, we found that each dynamical regime resembles a power-law growth (Figs. 4 and 6). The early-time dynamics was faster with a larger exponent.

Given the differences between early-time and late-time dynamics, any analysis that pools together data from different time points might produce inconsistent results that depend on the length of the experiment or simulation. This could partially explain the wide range of apparent power law exponents for boundary wandering and front roughness that are reported in the literature [17]. Similar issues arise in the analysis of the Kuramoto-Sivashinsky equation, which features growth instabilities related to those in our model [48–50]. Specifically, the Kuramoto-Sivashinsky equation exhibits a crossover between the EW dynamics at early times and KPZ dynamics at later times.

While our data is not sufficient to obtain reliable estimates of exponents or even validate the existence of a power law, we do observe clear deviations from the common theoretical models used to describe colony growth [17, 25, 42, 46]. At early times, we observe *β* that is inconsistent with the standard models and the boundary wandering is much faster than predicted by these models. At late times, the boundary wandering agrees with the standard models, but the value of *β* is too low. Thus, it appears that none of these models describes either early- or late-time dynamics correctly.

Our observations furthermore suggest that a thorough analysis of the power-law behavior should include both colony roughness and sector boundary displacements. Historically, however, most studies focused only one of these processes and, therefore, the agreement between observed and theoretically predicted exponents should be interpreted with some caution. Although we do not find an overwhelming support for any of the theoretical models, they do produce similar dynamics for some quantities some of the times, which could be exploited to speed up simulations. For example, one might adopt the Eden model to simulate the super-diffusive boundary wandering in the late-time regime.

Taken together, our findings provide a roadmap for rational control of morphological and genetic properties of microbial colonies. Upon combining dimensional analysis with simulations, one can predict the consequences of changing each of the biological or physical variables that affect metabolism, growth and motility. These predictions can then guide the selection of an optimal strategy to achieve a particular goal be it suppressing the evolution of drug resistance or slowing down the expansion rate. Given that nutrient limitation and cooperative motility play a major role in many cellular aggregates, we believe that our approach could be profitably deployed in a wider settings beyond microbial colonies.

## Acknowledgements

K.S.K. and A.G. were supported by the NIGMS through grant #1R01GM138530-01 and the Simons Foundation through grant #409704. I.D. and D.S. acknowledge funding from the Boston University Kilachand Funds as part of the Multicellular Design Program, from the NIH through grant R21CA260382, from the Human Frontiers Science Program (grant number RGP0060/2021), and from the U.S. Department of Energy, Office of Science, Office of Biological & Environmental Research through the Microbial Community Analysis and Functional Evaluation in Soils Science Focus Area Program (m-CAFEs) under contract number DE-AC02-05CH11231 to Lawrence Berkeley National Laboratory. Simulations were carried out on the Boston University Shared Computing Cluster.

